# Integrating Sample Similarities into Latent Class Analysis: A Tree-Structured Shrinkage Approach

**DOI:** 10.1101/2021.01.25.428033

**Authors:** Mengbing Li, Daniel E. Park, Maliha Aziz, Cindy M. Liu, Lance B. Price, Zhenke Wu

## Abstract

This paper is concerned with using multivariate binary observations to estimate the probabilities of unobserved classes with scientific meanings. We focus on the setting where additional information about sample similarities is available and represented by a rooted weighted tree. Every leaf in the given tree contains multiple samples. Shorter distances over the tree between the leaves indicate *a priori* higher similarity in class probability vectors. We propose a novel data integrative extension to classical latent class models (LCMs) with tree-structured shrinkage. The proposed approach enables 1) borrowing of information across leaves, 2) estimating data-driven leaf groups with distinct vectors of class probabilities, and 3) individual-level probabilistic class assignment given the observed multivariate binary measurements. We derive and implement a scalable posterior inference algorithm in a variational Bayes framework. Extensive simulations show more accurate estimation of class probabilities than alternatives that suboptimally use the additional sample similarity information. A zoonotic infectious disease application is used to illustrate the proposed approach. The paper concludes by a brief discussion on model limitations and extensions.

## 1. Introduction

### 1.1 Motivating Application

The fields of infectious disease epidemiology and microbial ecology need better tools for tracing the transmission of microbes between humans and other vertebrate animals (i.e., zoonotic transmissions), especially for colonizing opportunistic pathogens (COPs). Unlike frank zoonotic pathogens (e.g., *Salmonella*, SARS-CoV-2), the epidemiology of COPs, such as *Escherichia coli* (*E. coli*), *Staphylococcus aureus* (*S. aureus*) and *Enterococcus spp*., can be particularly cryptic due to their ability to asymptomatically colonize the human body for indefinite periods prior to initiating an infection, transmitting to another person, or being shed without a negative outcome (e.g., Price et al., 2017). Some COPs can colonize many different vertebrate hosts and cross-species transmissions can go unrecognized. Estimating the probability of zoonotic origin for a population of isolates and for each isolate would provide important insights into the natural history of infections and inform more effective intervention strategies, such as eliminating high-risk clones from livestock via vaccination.

Scientists often have two complementary sources of information: i) a phylogenetic tree constructed based on a few single nucleotide polymorphisms (SNPs) in the core genome shared by all isolates, where the leaves represent distinct core-genome multi-locus sequence types (STs, Maiden et al., 1998); the tree is useful for identifying a recent common ancestor for isolates that comprise an infectious disease outbreak; ii) presence or absence of multiple mobile genetic elements (MGE) that provide selective advantages in particular hosts and may be lost and gained as COPs transmit among hosts (e.g., Lindsay and Holden, 2004).

Recent research on two COP species, *E. coli* and *S. aureus*, has demonstrated the utility of complementing core-genome phylogenetic trees with host-associated MGEs to resolve host origins (e.g., Liu et al., 2018). However, in both cases only a single host-associated MGE was used. Analyses were largely limited to visual inspection of how each element fell on the scaffold of the evolutionary tree. For this approach to reach its full potential, we would need a statistical model that can 1) integrate phylogenetic information with the presence and absence of multiple host-associated MGEs, and 2) estimate the probability with which the isolates were derived from a particular host in each ST-specific population and for each individual isolate.

### 1.2 Integrating Sample Similarities into Latent Class Analysis

Based on multivariate binary data (e.g., presence or absence of multiple MGEs), we use latent class models (LCMs; e.g., Lazarsfeld, 1950; Goodman, 1974) to achieve the scientific goal of estimating the probabilities of unobserved host origins and perform individual-level probabilistic assignment of host origin. LCMs are examples of latent variable models that assume the observed dependence among multivariate discrete responses is induced by variation among unobserved or “latent” variables. It is well known that any multivariate discrete data distribution can be approximated arbitrarily closely by an LCM with a sufficiently large number of classes (Dunson and Xing, 2009, Corollary 1). The most commonly used LCMs assume the class membership indicators for the observations are drawn from a population with the same vector of class probabilities.

Trees or hierarchies are useful and intuitive for representing and reasoning about similarity or relation among objects in many real-world domains. We assume known entities at the leaves. In our context, each leaf may contain multiple observations or samples, each associated with the multivariate binary responses which are then combined to form the rows of a binary data matrix **Y**. In the motivating application, the latent class indicates the unobserved host origin (human or non-human) to be inferred by the presence or absence of multiple MGEs. The additional sample similarity information is represented by a maximum likelihood phylogenetic tree (e.g., Scornavacca et al., 2020). The leaves represent distinct contemporary core-genome *E. coli* STs.

To integrate tree-encoded sample similarity information into a latent class analysis, ad hoc groupings of the leaves may be adopted. From the finest to the coarsest leaf grouping, one may 1) analyze data from distinct lineages one at a time, 2) manually form groups of at least one leaf node and fit separate LCMs, or 3) fit all the data by a single LCM. However, all these methods pose significant statistical challenges. First, separate latent class analyses may have low accuracy in estimating latent class probabilities and other model parameters for rare lineages. Second, observations of similar lineages may have similar propensities in host jump resulting in similar host origin class probabilities. Modeling these similarities could lead to gain in statistical efficiency. Third, approaches based on coarse ad hoc groupings may obscure the study of the variation in the latent class probabilities across different parts of the tree. Finally, based on a single LCM or other approaches that use ad hoc leaf groupings, individual-specific posterior class probabilities can be averaged within in each leaf to produce a local estimate of the ***π***_*v*_. However, the ad hoc post-processing cannot fully address the issue of assessment of posterior uncertainty nor data-driven grouping of leaves, necessitating development of an integrative probabilistic modeling framework for uncertainty quantification and adaptive formation of leaf groups.

In this paper, we focus on integrating the tree-encoded sample similarity information into latent class analysis. We assume the tree information is given and not computed from the multivariate binary measurements. Observations in nearby leaves are assumed to have *a priori* similar propensities of being members of a particular class as characterized by the latent class probabilities. For example, higher similarities are indicated by shorter pairwise distances between observations. More generally, classical covariate-dependent latent class models (e.g., Bandeen-Roche et al., 1997; Formann, 1992) let the latent class probabilities vary explicitly as functions of observed covariates so that observations with more similar covariate values are assumed to have more similar latent class probabilities. Fully probabilistic tree-integrative methods have appeared in machine learning literature (e.g., Ghahramani et al., 2010; Roy et al., 2006; Ranganath et al., 2015) or in statistics for modeling hierarchical topic annotations (e.g., Airoldi and Bischof, 2016) or hierarchical outcome annotations based on given trees (e.g., Thomas et al., 2019). In epidemiology, Avila et al. (2014) proposed a two-stage approach to link patient clusters estimated from the tree and by the LCM results, which however remains ad hoc. However, current literature does not address probabilistic tree-integrative latent class analysis or adaptive formation of leaf groups for dimension reduction.

### 1.3 Primary Contributions

In this paper, we propose an unsupervised, tree-integrative LCM framework to 1) discover groups of leaves where multivariate binary measurements in distinct leaf groups have distinct vectors of latent class probabilities; And observations nested in any leaf group may belong to a pre-specified number of latent classes; 2) accurately estimate the latent class probabilities for each discovered leaf group and assign probabilities of an individual sample belonging to the latent classes; 3) leverage the relationship among the observations as encoded by the tree to boost the accuracy of the estimation of latent class probabilities. Without pre-specifying the leaf groups, the automatic data-driven approach enjoys robustness by avoiding potential mis-specification of the grouping structure. On the other hand, the discovered datadriven leaf groups dramatically reduce the dimension of leaves into fewer subgroups of leaves hence improving interpretation. In addition, the proposed approach shows better accuracy in estimating the latent class probabilities in terms of root mean squared errors, indicating the advantage of the shrinkage. On posterior computation, we derive a scalable inference algorithm based on variational inference (VI).

The rest of the paper is organized as follows. Section 2.2 defines tree-related terminologies and formulates LCMs. Section 3 proposes the prior for tree-structured shrinkage in LCMs. Section 4 derives a variational Bayes algorithm for inference. Section 5 compares the performances of the proposed and alternative approaches via simulations. Section 6 illustrates the approach by analyzing an *E. coli* data set. The paper concludes with a brief discussion.

## 2. Model

We first introduce necessary terminologies and notations to describe a rooted weighted tree. LCMs are then formulated for data on the leaves of the tree.

### 2.1 Rooted Weighted Trees

A rooted tree is a graph 𝒯 = (𝒱, *E*) with node set 𝒱 and edge set *E* where there is a root *u*_0_ and each node has at most one parent node. Let *p* = |𝒱| represent the total number of leaf and non-leaf nodes. Let 𝒱_*L*_ ⊂ 𝒱 be the set of leaves, and *p*_*L*_ = |𝒱_*L*_| *< p*. We typically use *u* to denote any node (*u* ∈ 𝒱) and *v* to denote any leaf (*v* ∈ 𝒱_*L*_). Each edge in a rooted tree defines a *clade*: the group of leaves below it. Splitting the tree at an edge creates a partition of the leaves into two groups. For any node *u* ∈ 𝒱, the following notations apply: *c*(*u*) is the set of offspring of *u, pa*(*u*) is the parent of *u, d*(*u*) is the set of descendants of *u* including *u*, and *a*(*u*) is the set of ancestors of *u* including *u*. In Figure 3(a), if *u* = 2, then *c*(*u*) = {6, 7, 8}, *pa*(*u*) = {1}, *d*(*u*) = {2, 6, 7, 8}, and *a*(*u*) = {1, 2}. The phylogenetic tree in our motivating application is a nested hierarchy of 133 STs for *N* = 2, 663 observations, where the *p*_*L*_ = |𝒱_*L*_| = 133 leaves represent distinct STs and the *p* − *p*_*L*_ = 132 internal (non-leaf) nodes represent ancestral *E. coli* strains leading up to the observed leaf descendants.

Edge-weighted graphs appear as a model for numerous problems where nodes are linked with edges of different weights. In particular, the edges in 𝒯 are attached with weights where *w* : *E* → ℝ^+^ is a weight function. Let 𝒯_*w*_ = (𝒯, *w*) be a rooted weighted tree. A path in a graph is a sequence of edges which joins a sequence of distinct vertices. For a path *P* in the tree connecting two nodes, *w*(*P*) is defined as the sum of all the edge weights along the path, often referred to as the “length” of *P*. The distance between two vertices *u* and *u*^′^, denoted by 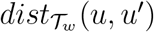 is the length of a shortest (with minimum length) (*u, u*^′^)-path. 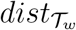 is a distance: it is symmetric and satisfies the triangle inequality. In our motivating application, the edge length represents the number of nucleotide substitutions per position; the distance between two nodes provides a measurement of the similarity or divergence between any two core-genome sequences of the input set. In this paper, we use *w*_*u*_ to represent the edge length between a node *u* and its parent node *pa*(*u*). *w*_*u*_ is fully determined by 𝒯_*w*_. For the root *u*_0_, there are no parents, i.e. *pa*(*u*_0_) = ∅; we set 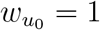.

### 2.2 Latent Class Models for Data on the Leaves

Although LCMs can deal with multiple categorical responses in general, for simpler presentations in this paper, we focus on presenting the model and algorithm using multivariate binary responses and their application to the motivating data.

#### Notations

Let 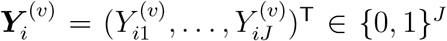 be the vector of binary responses for observation *i* ∈ [*n*_*v*_] that is nested within leaf node *v* ∈ 𝒱_*L*_, where *n*_*v*_ is the number of observations in leaf *v*. Throughout this paper, let [*Q*] := {1, …, *Q*} denote the set of positive integers smaller than or equal to *Q*, where *Q* is a positive integer. Let 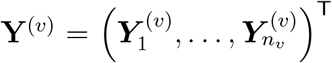 be the data from observations in leaf *v*. Let **Y** = ((**Y**^(1)^)^T^, …, (**Y**^(*pL*)^)^T^)^T^ represent the binary data matrix with *N* = ∑v∈𝒱_*L*_ *n* _*v*_ rows and *J* columns. Let ℒ = (*v*_1_, …, *v*_*N*_)^T^ be the “sample-to-leaf indicators” that map every row of data **Y** into a leaf in 𝒯_*w*_. Sample similarities are then characterized by between-leaf distances in 𝒯_*w*_. In this paper, we assume ℒ and 𝒯_*w*_ are given and focus on incorporating (ℒ, 𝒯_*w*_) into a statistical model for **Y**.

#### LCM for Data on the Leaves

The LCM is specified in two steps:

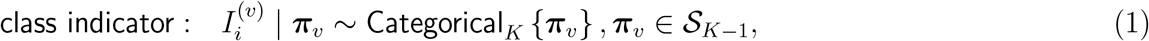

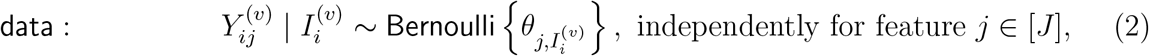

and independently for observation *i* ∈ [*n*_*v*_] and leaf node *v* ∈ 𝒱_*L*_. Here *K* is a pre-specified number of latent classes in the context of the application, e.g., *K* = 2 for unobserved human and non-human hosts; see Section 4.1 for a simple strategy in applications where data-driven *K* is desired. In addition, 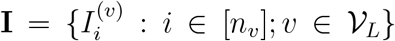 represent the latent class indicators and 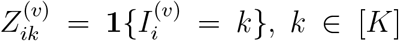, where **1**{*A*} is an indicator function which equals 1 if statement *A* is true and 0 otherwise; Let 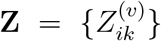. We have assumed observations in different leaves have potentially different vectors of class probabilities ***π***_*v*_ = (*π*_*v*1_, …, *π*_*vK*_)^T^ ∈ 𝒮_*K*−1_, *v* ∈ 𝒱_*L*_, where 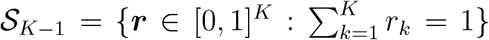 is the probability simplex. *θ*_*jk*_ ∈ [0, 1] is the positive response probability for feature *j* ∈ [*J* ] in class *k* ∈ [*K*]. In our motivating application, the MGEs adapt to the unobserved type of host origin (i.e., latent class) which can be characterized by class-specific response probability profiles ***θ***_·*k*_ = (*θ*_1*k*_, …, *θ*_*Jk*_)^T^, *k* ∈ [*K*]; let **Θ** = (***θ***_·1_, …, ***θ***_·*K*_)^T^. Because the latent class indicators 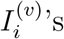 are assumed to be unobserved, the observed data likelihood for *N* observations is 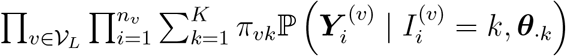.

Throughout this paper, we assume that we wish to classify individuals into *K* classes with the same set of (***θ***_·1_, …, ***θ***_·*K*_) so classes have coherent interpretation. However, we do not assume that observations are drawn from a population with a single vector of latent class probabilities. Figure 1 provides a schematic of the data generating mechanism given ***π***_*v*_ for three leaves.

**Figure 1:**
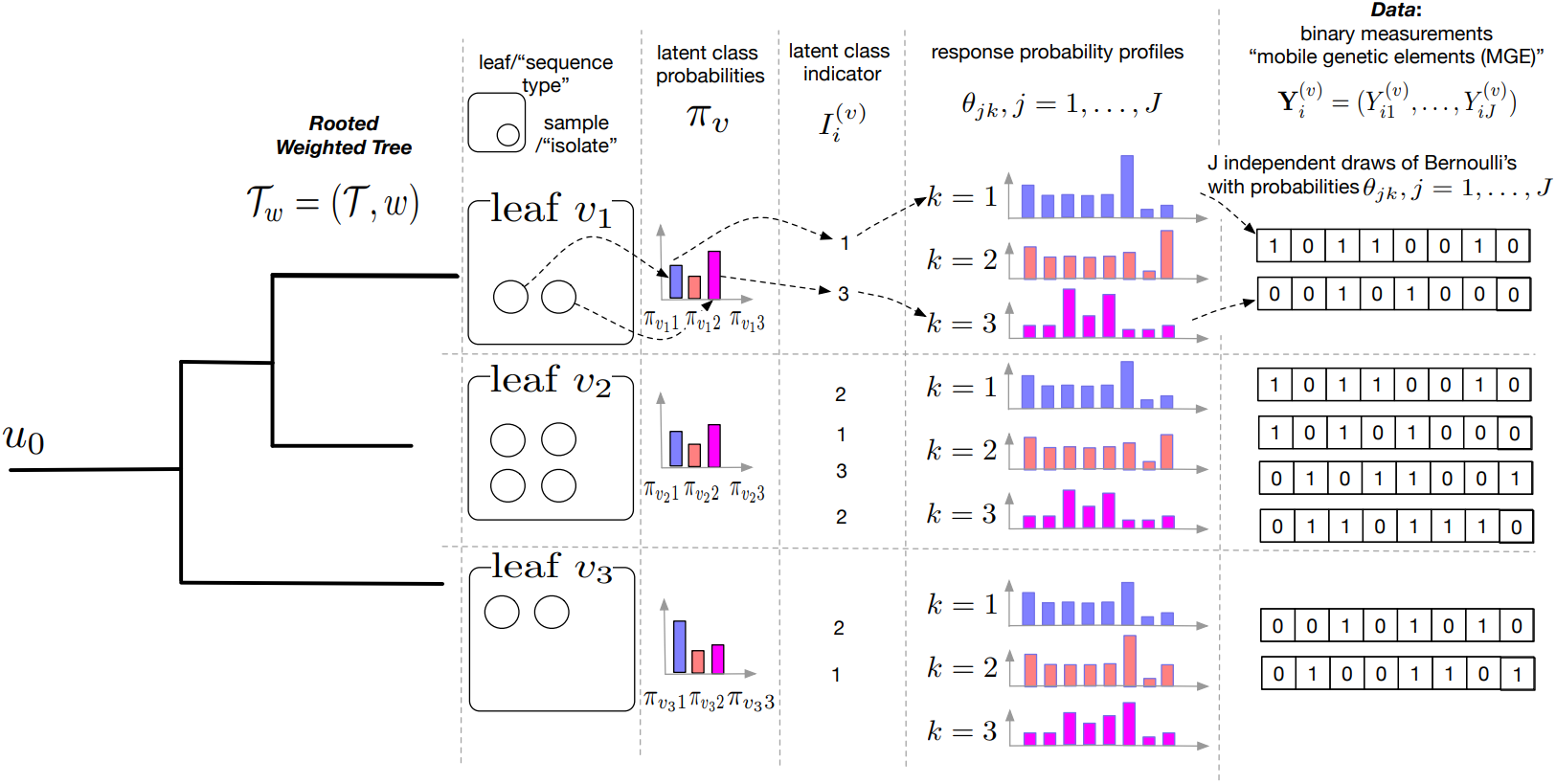
Schematic representation of a hypothetical rooted weighted tree with three leaves and data generated based on the proposed model with *K* = 3 latent classes, 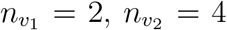 and 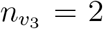, *J* = 8. This figure appears in color in the electronic version of this article, and any mention of color refers to that version

## 3. Prior Distribution

We first specify a prior distribution for {***π***_*v*_ : *v* ∈ 𝒱_*L*_}. Because leaf-specific sample sizes may vary, we propose a tree-structured prior to borrow information across nearby leaves. The prior encourages collapsing certain parts of the tree so that observations within a collapsed leaf group share the same vector of latent class probabilities. In particular, we extend Thomas et al. (2019) to deal with rooted weighted trees in an LCM setting. The prior specification is completed by priors for the class-specific response probabilities **Θ**.

### Tree-structured prior for latent class probabilities **π**_*v*_

We specify a spike-and-slab Gaussian diffusion process prior along a rooted weighted tree based on a logistic stick-breaking parameterization of ***π***_*v*_. We first reparameterize ***π***_*v*_ with a stick-breaking representation: *π*_*vk*_ = *𝒱*_*vk*_ Π_*s<k*_(1 − *𝒱*_*vs*_), for *k* ∈ [*K*], where 0 ⩽ *𝒱*_*vk*_ ⩽ 1, for *k* ∈ [*K* − 1] and *𝒱*_*vK*_ = 1.

We further logit-transform *𝒱*_*vk*_, *k* ∈ [*K* − 1], to facilitate the specification of a Gaussian diffusion process prior without range constraints. In particular, let *η*_*vk*_ = *σ*^−1^(*𝒱*_*vk*_), *k* ∈ [*K* − 1], *v* ∈ 𝒱_*L*_, where *σ*(*x*) = 1*/*{1 + exp(−*x*)} is the sigmoid function. The logistic stick-breaking parameterization is completed by

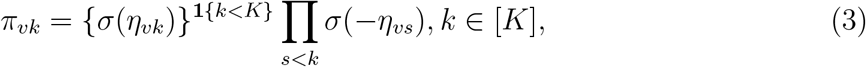

which affords simple and accurate posterior inference via variational Bayes (see Section 4). For a leaf *v* ∈ 𝒱_*L*_, let

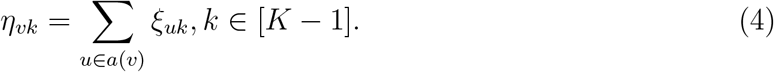

Here *η*_*vk*_ is defined for leaves only and *ξ*_*uk*_ is defined for all the nodes. Suppose *v* and *v*^′^ are leaves and siblings in the tree such that *pa*(*v*) = *pa*(*v*^′^), setting *ξ*_*vk*_ = *ξ*_*v ′k*_ = 0 implies *η*_*vk*_ = *η*_*v k*_ for *k* ∈ [*K* − 1], and hence ***π***_*v*_ = ***π***_*v*_. More generally, a sufficient condition for *M* leaves *η*_*vk*_, *v* ∈ {*v*_1_, …, *v*_*M*_} to fuse is to set *ξ*_*uk*_ = 0 for any *u* that is an ancestor of any of {*v*_1_, …, *v*_*M*_} but not common ancestors for all *v*_*m*_. That is, to achieve grouping of observations that share the same vector of latent class probabilities, in our model, it is equivalent to parameter fusing. In the following, we specify a prior on the *ξ*_*uk*_ that *a priori* encourages sparsity, so that closely related observations are likely grouped to have the same vector of class probabilities. The fewer distinct ancestors two nodes have, the more likely the parameters *η*_*vk*_ are fused, because the prior would encourage fewer auxiliary variables *ξ*_*uk*_ to be set to zero. In particular, we specify

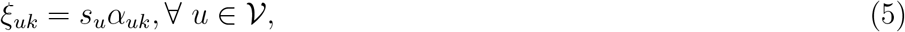

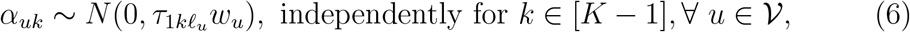

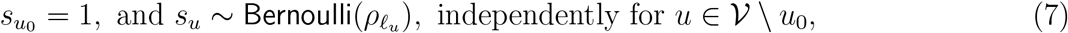

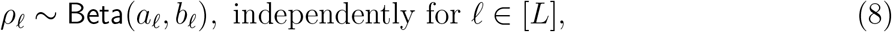

where *N* (*m, s*) represents a Gaussian density function with mean *m* and variance *s. τ*_1*k*_ is the unit-length variance and controls the degree of diffusion along the tree which may differ by dimension *k* and node level *£*_*u*_ where *£*_*u*_ ∈ [*L*] represents the “level” or “hyperparameter set indicator” for node *u*. For example, in simulations and data analysis, we will assume that the root for the diffusion process has a prior unit-length variance distinct from other non-root nodes. For the root *u*_0_ with 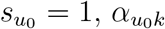 initializes the diffusion of *η*_*uk*_.

Leaf groups are formed by selecting a subset of nodes in 𝒱: 𝒰 = {*u* ∈ 𝒱 : *s*_*u*_ = 1}. Except a probability-zero set, two leaves *v* and *v*^′^ are grouped, or “fused”, if and only if *a*(*v*) ∩ 𝒰 = *a*(*v*^′^) ∩ 𝒰. In particular, the null set is {*η*_*vk*_ = *η*_*vk′*_, *k* ∈ [*K* − 1]} ∩ {Σ_*u*∈[*a*(*v′*)∩U]\[*a*(*v*)∩U]_ *α*_*uk*_ = Σ _*u*∈[*a*(*v*)∩U]\[*a*(*v′*)∩U]_ *α*_*uk*_} where the latter has probability zero. In Section 4.1, we will estimate 𝒰 using the posterior median model.

#### Remark 1

Equations (4)-(8) define a Gaussian diffusion process initiated at 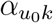 :

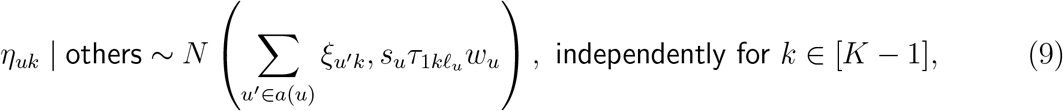

for any non-root node *u* ≠ *u*_0_; also see the seminal formulation by Felsenstein (1985). To aid the understanding of this Gaussian diffusion prior, it is helpful to consider a special case of *s*_*u*_ = 1 and *£*_*u*_ = 1, ∀*u* ∈ 𝒱. For two leaves *v, v*^′^ ∈ 𝒱_*L*_, the prior correlation between *η*_*vk*_ and *η*_*v′ k*_ is

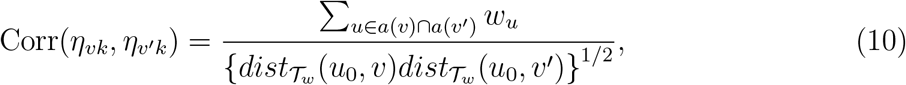

When *v* and *v*^′^ have the same number of ancesters (|*a*(*v*)| = |*a*(*v*^′^)|) and all edges have identical weight *w*_*u*_ = *c*, ∀*u*, the prior correlation is the fraction of common ancestors. Note that ***η***_*v*_ fully determines ***π***_*v*_ in (3) and induces correlations among {***π***_*v*_, *v* ∈ V_*L*_}.

#### Remark 2

One reviewer raised an important question on the choice of encouraging prior correlation among {***π***_*v*_} rather than among the latent class indicators 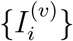. In the present prior distribution, by integrating out {***π***_*v*_}, we have induced prior marginal correlation among 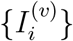 for observation in nearby leaves. Additional prior correlation amongst the 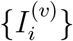 can be introduced via an additional layer of prior over the 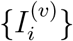 conditional on {***π***_*v*_}, e.g., through clustered samples. The absence of such clustered sampling structure in the motivating application points us towards the former simpler strategy.

### Priors for class-specific response probabilities

Let *γ*_*jk*_ = log {*θ*_*jk*_*/*(1 − *θ*_*jk*_)}. We specify

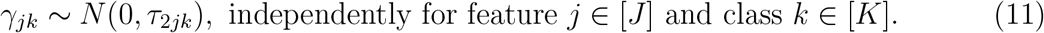

### Joint distribution

Let ***β*** = (**Z, *s, γ, α, g***) collect all the unknown parameters where ***s*** = {*s*_*u*_ : *u* ∈ V}, ***γ*** = {*γ*_*jk*_, *j* ∈ [*J* ]; *k* ∈ [*K*]}, ***α*** = {*α*_*uk*_ : *u* ∈ V, *k* ∈ [*K* − 1]}, ***g*** = (*ρ*_1_, …, *ρ*_*L*_)^T^, ***a*** = (*a*_1_, …, *a*_*L*_)^T^, and ***b*** = (*b*_1_, …, *b*_*L*_)^T^. Hereafter we use pr(*A* | *B*) to denote a probability density or mass function of quantities in *A* with parameters *B*; when *B* represents hyperparameters or given information in this paper, we simply use pr(*A*), e.g., we will use pr(**Y, *β***) to represent pr(**Y, *β*** | ***τ***_1_, ***τ***_2_, ***a, b***, 𝒯_*w*_, ℒ). The joint distribution of data and unknown quantities can thus be written as:

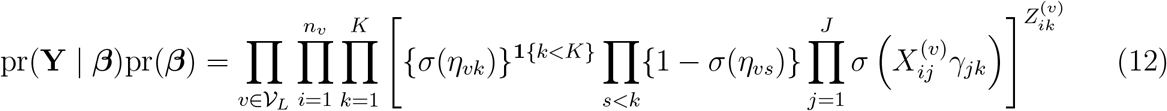

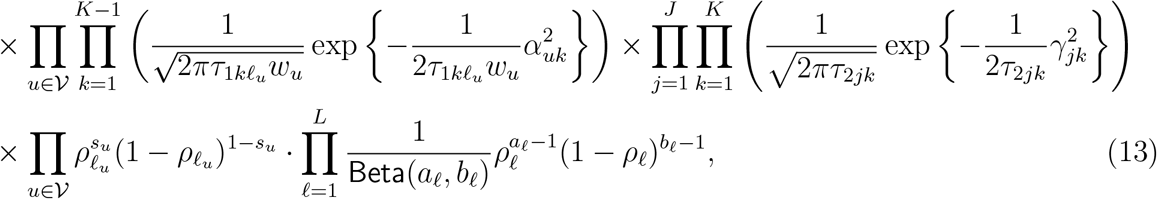

where 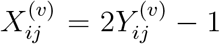. Tree information 𝒯_*w*_ enters the joint distribution in the definition of ***η***_*v*_ (Equations (4)); sample-to-leaf indicators ℒ choose among {***η***_*v*_, *v* ∈ 𝒱_*L*_} for every observation in Equation (12). By setting *s*_*u*_ = 0 for all the non-root nodes in Equation (5), the classical LCM with a single 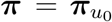 results. Figure 2 shows a directed acyclic graph (DAG) that represents the model likelihood and prior specifications.

**Figure 2:**
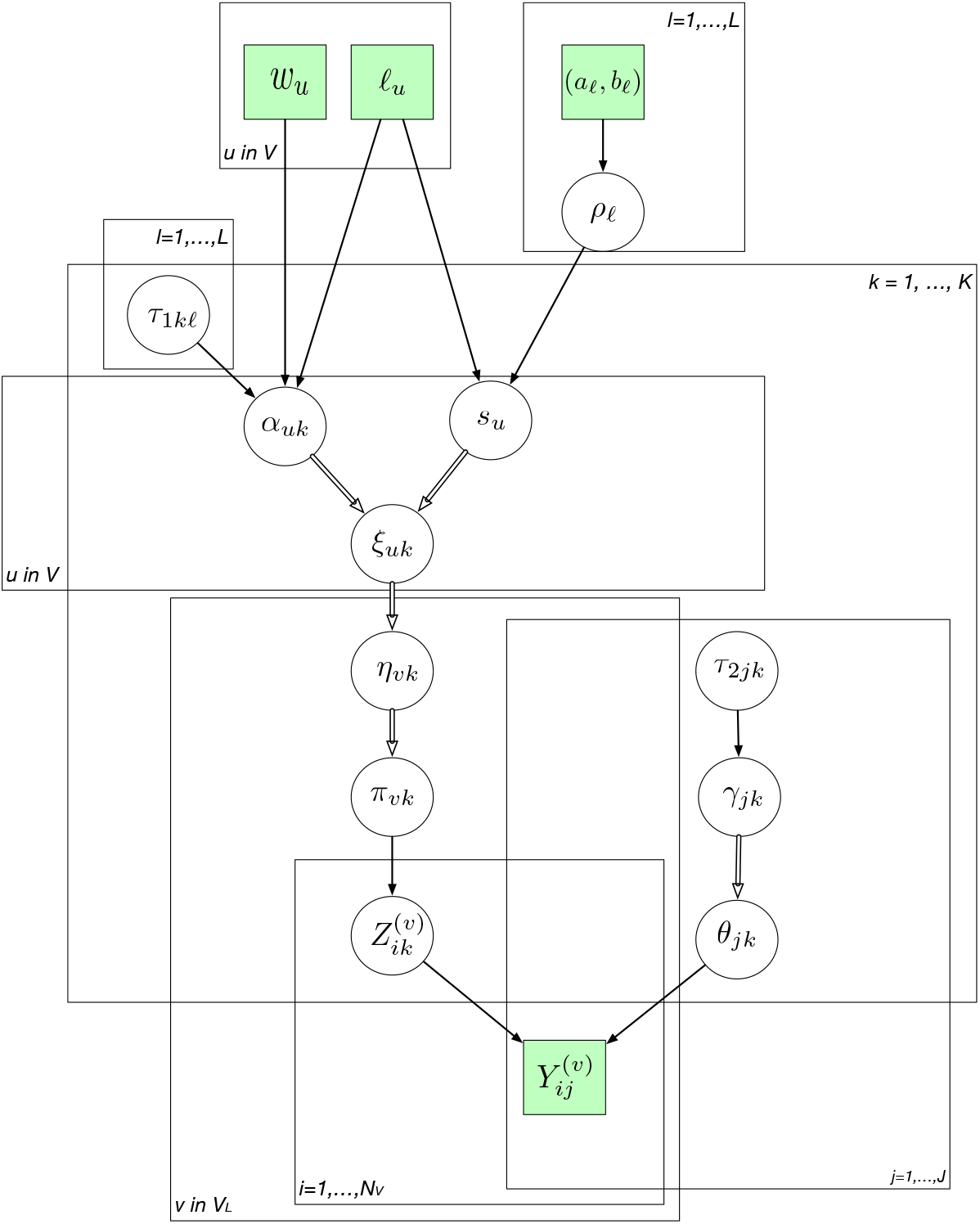
The directed acyclic graph (DAG) representing the structure of the model likelihood and priors. The quantities in squares are either data or hyperparameters; the unknown quantities are shown in the circles. The arrows connecting variables indicate that the parent parameterizes the distribution of the child node (solid lines) or completely determines the value of the child node (double-stroke arrows). The rectangular “plates” where the variables are enclosed indicate that a similar graphical structure is repeated over the index; The index in a plate indicate nodes, hyperparameter levels, leaves, subjects, classes and features. This figure appears in color in the electronic version of this article, and any mention of color refers to that version.

**Figure 3:**
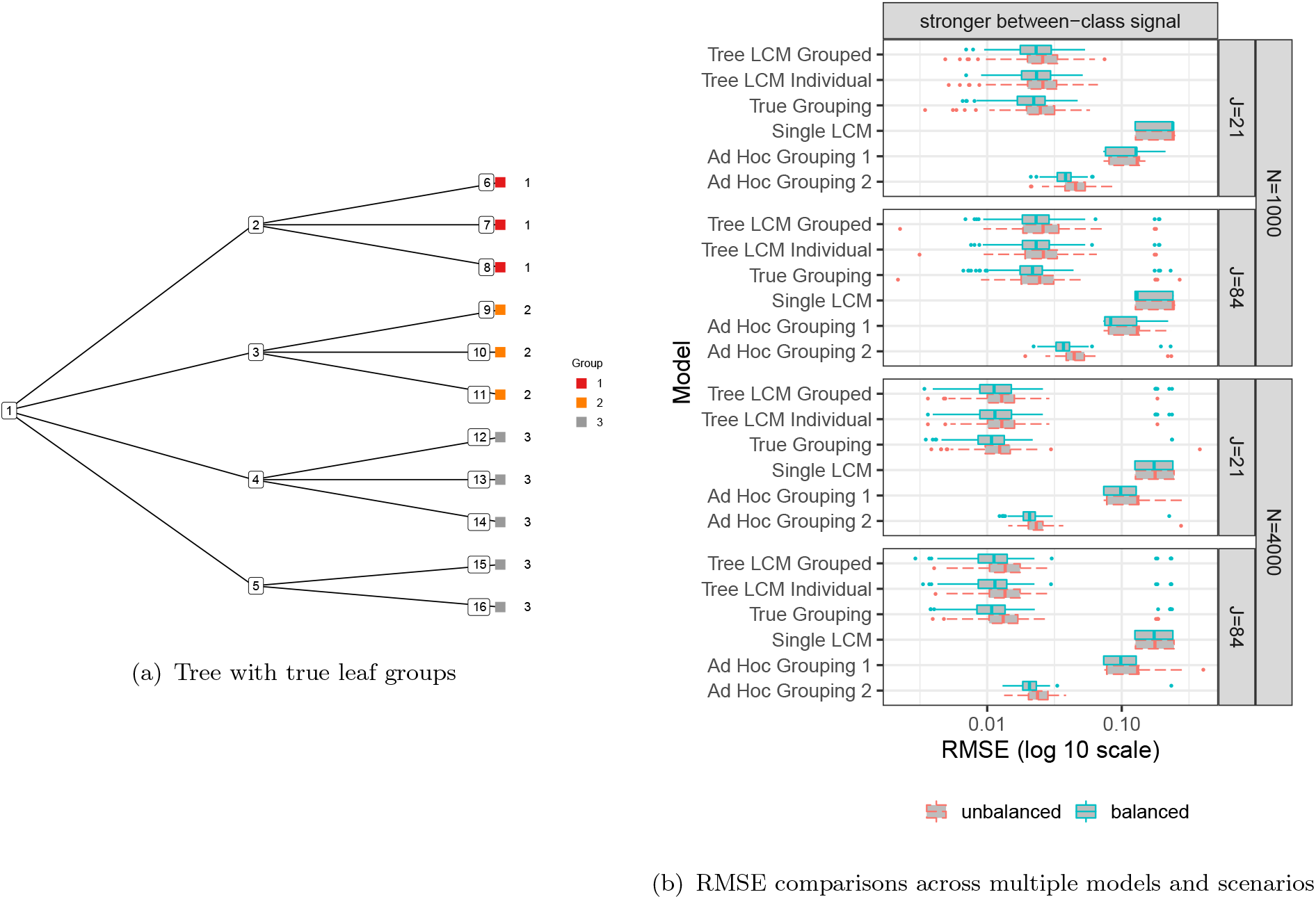
Simulation studies show the proposed model produces grouped estimates 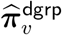 with similar or smaller RMSEs compared to alternatives (see Section 5). This figure appears in color in the electronic version of this article, and any mention of color refers to that version.

## 4. Variational Inference Algorithm

Calculating a posterior distribution often involves intractable high-dimensional integration over the unknowns in the model. Traditional sequential sampling approaches such as Markov chain Monte Carlo (MCMC) remains a widely used inferential tool based on approximate samples from the posterior distribution. They can be powerful in evaluating multidimensional integrals. However, they do not guarantee closed-form posterior distributions. Variational inference (VI) is a popular alternative to MCMC for approximating the posterior distribution and has been widely used in machine learning and gaining interest in statistics (e.g., Blei et al., 2017; Ormerod and Wand, 2010). In particular, VI has also been used for fitting the classical LCMs (e.g., Grimmer, 2011). VI requires a user-specified family of distributions that can be expressed in tractable forms while being flexible enough to approximate the true posterior; the approximating distributions and their parameters are referred to as “variational distributions” and “variational parameters”, respectively. VI algorithms find the best variational distribution that minimizes the Kullback-Leibler (KL) distance between the variational family and the true posterior distribution. VI has been widely applied in Gaussian (Carbonetto and Stephens, 2012; Titsias and Lázaro-Gredilla, 2011) and binary likelihoods (e.g., Jaakkola and Jordan, 2000; Thomas et al., 2019). Also see Blei et al. (2017) for a detailed review. We use VI because it is fast, bypasses infeasible analytic integration or data augmentation that is otherwise needed for MCMC under Dirac spike components and prior-likelihood non-conjugacy (Tüchler, 2008), and enables data-driven selection of hyperparameters via approximate empirical Bayes (Equation (S8), Supporting Information) at the cost of slight variance-covariance under-estimation, the degree of which we assess in Section 5.

We use VI algorithm to conduct inference using variational distributions factorized as:

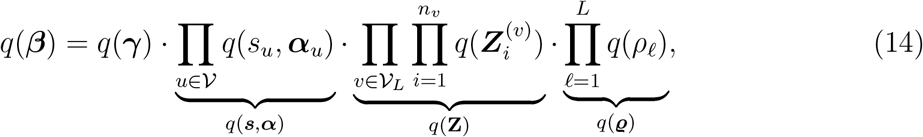

where 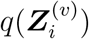 is a multinomial distribution with variational parameters 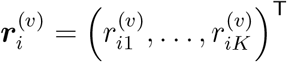, and 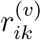 represents the approximate posterior probability of observation *i* in leaf *v* belonging to class *k* and 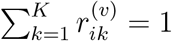. Importantly, we make no other assumptions about the particular parametric form of variational distributions, which by the VI updating rules can be shown to take familiar distributional forms (see Appendix A).

VI finds *q* that minimizes the Kullback-Leibler (KL) distance between the variational family and the true posterior distribution: 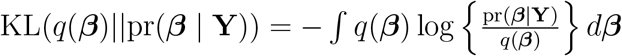. However, the KL distance depends on the intractable posterior distribution is not easily computed. Fortunately, based on a well-known equality log pr(**Y**) = ε(*q*) + KL(*q*(***β***) ‖ pr(***β*** | **Y**)), where 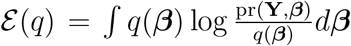 is referred to as evidence lower bound (ELBO) because log pr(**Y**) ⩾ ε(*q*). Because pr(**Y**) is a constant, minimizing the KL divergence is equivalent to maximizing E(*q*). The VI algorithm updates each component of *q*(***β***) in turn while holding other components fixed. However, because of the nonlinear sigmoid functions in Equation (12), generic VI updating algorithms for *q*(*s*_*u*_, ***α***_*u*_) and *q*(***γ***) involve integrating over random variables in the sigmoid function hence lack closed forms. To make the updates analytically tractable, we replace Equation (12) with an analytically tractable lower bound. In particular, we use a technique introduced by Jaakkola and Jordan (2000) which bounds the sigmoid function from below by a Gaussian kernel with a tuning parameter, hence affords closed-form VI updates; also see Durante et al. (2019) for a modern view of this technique as a bona fide mean-field approximation with Pòlya Gamma data augmentation. In particular, we will use the inequality

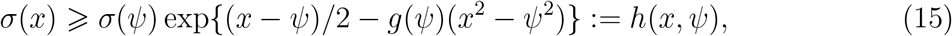

with 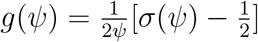 where *ψ* is a tuning parameter.

We approximate ELBO *ε*(*q*) by *ε*^*^(*q*):

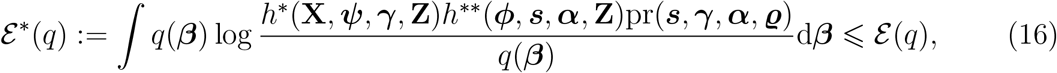

where 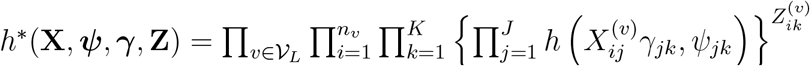, and 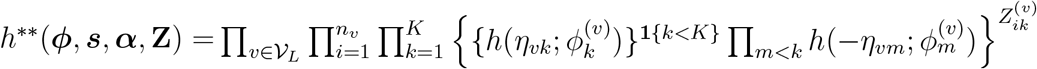. The VI algorithm iterates until convergence to find the optimal variational distribution *q* that maximizes ε^*^(*q*). Because ε^*^(*q*) log *π*(**Y**), it can be viewed as an approximation to the marginal likelihood. We maximize over ***ψ*** and ***ϕ*** to obtain the best approximation. In addition, we adopt an approximate empirical Bayes approach by optimizing the VI objective function ε^*^(*q*) over the hyperparameters ***τ***_1_ and ***τ***_2_. Relative to specifying weakly informative but often non-conjugate hyperprior for the variance parameters, optimizing hyperparameter is more practically convenient (e.g., Thomas et al., 2019). Because updating the hyperparameters changes the prior, we need to update *q*, ***ψ*** and ***ϕ*** again. This leads to an algorithm that alternates between maximizing ε^*^(*q*) in (*q*, ***ψ, ϕ***) and in (***τ***_1_, ***τ***_2_) until convergence. We update the hyperparameters every *d* complete VI iterations. Pseudocode in Algorithm 1 outlines the VI updates; Appendix A1 details the exact updating formula.

### 4.1 Posterior Summaries

Two sets of point and interval estimates for {***π***_*v*_ : *v* ∈ 𝒱_*L*_} are available from the VI algorithm: 1) data-driven grouped (“fused”) estimates 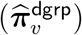 that are formed by setting a subset of ***s*** to one and the rest to zero, and 2) leaf-specific estimates 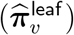. For 1), we select the posterior median model by setting *s*_*u*_ = 1 for nodes in 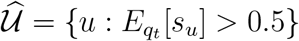 (see Step 1b, Appendix A1). For leaves *v* and *v*^′^, 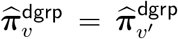 if and only if 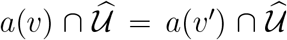. Because no closed-form posterior distributions for ***π***_*v*_ are readily available under logistic stick-breaking representation, we compute the approximate posterior mean and approximate 95% credible intervals (CrIs) by a Monte Carlo procedure after convergence of Algorithm 1. For 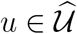, we first draw *B* = 10^5^ random independent samples of *α*_*uk*_ from *N* (*E*_*q*_ [*α*_*uk*_ | *s*_*u*_ = 1], V_*gt*_ [*α*_*uk*_ | *s*_*u*_ = 1]), for *k* ∈ [*K* − 1]. We then compute *B* corresponding ***π***_*v*_ vectors based on Equations (3) to (5) with 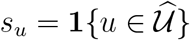 in (5). Finally, we compute the empirical means and 95% CrIs marginally for *π*_*vk*_, *k* ∈ [*K*]. The above Monte Carlo procedure is extremely fast given only independent Gaussian samples are drawn. As a comparison, for 2), we define leaf-specific estimates 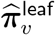 by the mean of (3) where 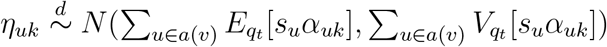, for *k* ∈ [*K*]. We also use Monte Carlo simulation to approximate the posterior means and 95% CrIs. In general, 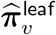 differ across the leaves. In contrast, the data-driven grouped estimates 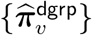 induce dimension reduction.

### Prediction

The out-of-sample predictive probability of class *k* for a new observation nested in leaf *v* is 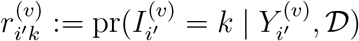, where D = (**Y**, 𝒯_w_, ℒ, ***a, b, τ***_1_, ***τ***_2_). We have

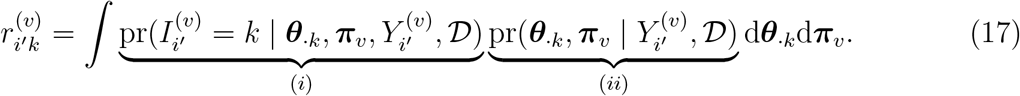

We approximate (17) by plug-in estimators: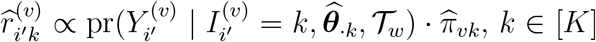. This can be seen by noting that term 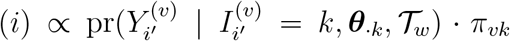, and term (*ii*) ≈ pr(***θ***_·*k*_, ***π***_*v*_ | 𝒟) which we approximate by a Dirac measure at 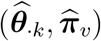. Here 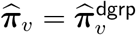.

### Choice of K

In applications where data-driven selection of *K* is more desirable, we may follow Bishop (2006) and use criterion 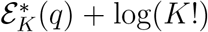 where 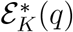 is the lower bound of log marginal data likelihood for a *K*-class model and the correction term is to make different models comparable (e.g., Grimmer, 2011, Section 5.2).

## 5. Simulation

### 5.1 Design and Performance Metrics

We conducted a simulation study to evaluate the performance of the proposed tree-integrative LCM. We compare our model to a few alternatives with ad hoc grouping of observations in terms of accuracy in estimating {***π***_*v*_, *v* ∈ *V*_*L*_}. Data were generated under two scenarios with different class-specific response profiles **Θ**. Appendix A2 details the true parameter settings of the simulations. Figure 3(a) visualizes the tree 𝒯_*w*_ with equal edge weights and true leaf groups used in the simulation with *p*_*L*_ = 11 leaves and *G* = 3 groups.

We simulated *R* = 200 independent replicate data sets for different total sample sizes (*N* = 1000, 4000). For each *N*, we set *n*_*v*_ ≈ *N/p*_*L*_ for *ν* ∈ 𝒱_*L*_ (with rounding where needed) to investigate balanced leaves and set *n*_*v*_ to be approximately 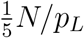 or 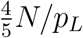 with equal chance for mimicking unbalanced observations across leaves. For observations in a leaf *v*, we simulate 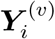 according to an LCM with class probabilities ***π***_*v*_ and class-specific response probabilities **Θ**. We simulated data for different dimensions *J* = 21, 84, for *K* = 3 classes.

For each simulated data set, we fitted the proposed model, based on which we compute 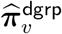 and 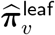 (see Section 4.1). Our primary interest is in 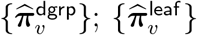 are for comparisons. In addition, we also tested a few approaches based on ad hoc leaf node groupings: 1) True grouping analysis (fit separate LCMs to obtain estimates in each of the true groups); 2) Single group LCM analysis (omit sample-to-leaf indicators ℒ, hence the tree information);3) Ad hoc grouping 1 (manual grouping coarser than the true grouping); 4) Ad hoc grouping 2: classical LCMs for data on each leaf. All analyses assume **Θ** does not vary by leaves.

We used three model performance metrics. First, we computed the root mean squared errors (RMSE) for an estimate 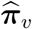 where 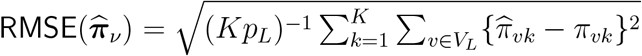 Second, we compared the true and the estimated leaf groupings via adjusted Rand Index (ARI, Hubert and Arabie, 1985). ARI is a chance-corrected index that takes value between −1 and 1 with values closer to 1 indicating better agreement. Finally, we estimated the coverage probability of the approximate 95% CrIs. For each true group *g*, we compute the frequency of the approximate 95% CrI (computed along with 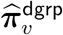) containing the truth, conditional on the event that an estimated partition of the leaf nodes includes *g*.

### 5.2 Simulation Results

Figure 3 shows comparisons among the RMSEs for different models under different scenarios. For sample sizes *N* = 1000 and *N* = 4000, the proposed methods with data-driven grouping 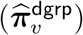 produced similar or better RMSE than analyses based on ad hoc leaf groupings, which restrict leaves into incorrect groupings that are coarser (single LCM and ad hoc grouping 1) or finer (ad hoc grouping 2) than the truth. The proposed approach 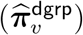 achieved similar RMSE as 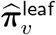, indicating little accuracy was lost in exchange for dimension reduction. The RMSEs of 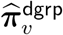 were similar to estimates of ***π***_*v*_, *v* ∈ 𝒱_*L*_ obtained from analyses based on the true leaf grouping. Indeed, the accuracy of group discovery increased with sample sizes with other settings fixed. Average ARIs across replications for each scenario were high (0.94 to 0.99) indicating good recovery of the true leaf groups. Although the groups discovered were not perfect, the comparable RMSEs suggest desirable adaptability of the proposed approach in effective collapsing of the leaves. The RMSE for 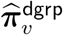 was smaller than analyses based on a refined leaf-level grouping: smaller sample sizes in the leaves resulted in loss of efficiency in separate estimations of ***π***_*v*_ across leaves. RMSEs were further reduced under a larger *J* or balanced sample sizes in the leaves. However, we again observed similar relative advantage of the proposed 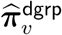. The relative comparisons of RMSEs under less discrepant true class-specific response profiles remained similar (see Appendix Figure S2).

The observed coverage rates of the approximate 95% CrIs achieved the nominal level satisfactorily (see Appendix Figure S1). Slight under-coverage occurred under smaller *N*, unbalanced sample sizes, smaller *J* and leaf groups with smaller number of observations.

This is partially a consequence of VI as an inner approximation to the posterior distribution which may underestimate the posterior uncertainty (e.g., Chapter 10, Bishop, 2006).

Finally, we also considered scenarios where only a single group of leaves is present in truth for which the classical LCM is perfectly appropriate. Appendix Figure S3 shows, by learning the posterior node-specific slab-versus-spike selection probabilities, the proposed model produces similar RMSEs as the classical LCM.

## 6. E. Coli Data Application

### 6.1 Background and Data

*E. coli* infections cause millions of urinary tract infections (UTIs) in the US each year (e.g., Johnson and Russo, 2002). Many studies have shown that extraintestinal pathogenic *E. coli* (ExPEC) strains routinely colonize food animals and contaminate the food supply chain serving as a likely link between food-animal *E. coli* and human UTIs (e.g., Johnson et al., 2005). The scientific team adopted a novel strategy of augmenting fine-scale core-genome phylogenetics with interrogation of accessory host-adaptive MGEs (see Section 1.1). The scientific goal is to accurately estimate the probabilities of *E. coli* isolates with human and non-human host-origins across genetically diverse but related *E. coli* sequence types (STs). We restrict our analysis to *N* = 2, 663 *E. coli* isolates in a well-defined collection from humans and retail meat obtained over a 12-month period in Flagstaff, Arizona, US. Each isolate belongs to one of *p*_*L*_ = 133 different STs (leaves in the phylogenetic tree) that are identified via a multilocus sequence typing scheme based on short-read DNA sequencing. A total of *J* = 17 MGEs were curated and associated with functional annotations. Each ST was represented by at least four isolates. We constructed rooted, maximum-likelihood phylogenies using core-genome SNP data for the 133 STs. Figure 4 shows the estimated phylogenetic tree for the STs where the edge lengths represents the substitution rate in the conserved core genome. Every ST is overlaid in the same row with the empirical frequencies of 1) *J* = 17 MGEs and 2) the observed sources (human clinical or meat samples) which may differ from the true host origin. The observed frequencies of the MGEs vary greatly across lineages. We apply the proposed tree-integrative LCM to 1) estimate the probabilities of *unobserved* human and non-human host-origins for all *E. coli* STs with data-driven groupings of the STs for dimension reduction; and 2) to produce isolate-level probabilistic host-origin assignment. The context of the study restricts us to assume the host origin of each isolate is in one of two unobserved class of human vs food animals. A subset of preliminary data is analyzed in this paper for illustrating the proposed method. Inclusion of additional samples and/or MGEs may change findings. The final results and the detailed workflow of MGE discovery will be reported elsewhere.

**Figure 4:**
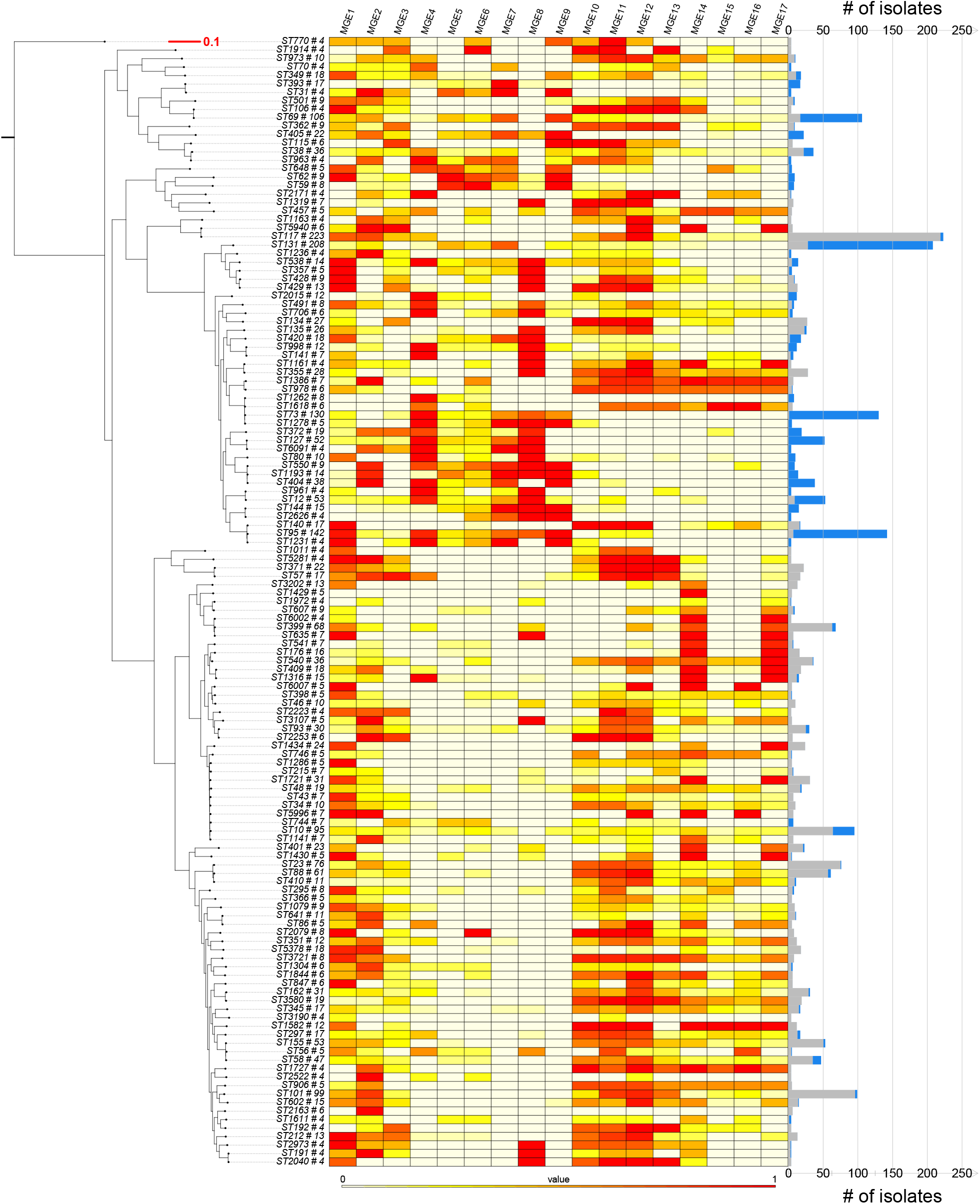
The empirical frequencies for *J* = 17 MGEs within each ST mapped in the core-genome phylogenetic tree. The red scale bar represents the substitution rate in the conserved core genome. The bars on the right indicate the total number isolates of each ST; the gray and blue bars represent the number of isolates obtained from apparent non-human and human sources, respectively. The core-genome phylogenetic tree on the left margin maps *N* = 2, 663 *E. coli* isolates into *p*_*L*_ = 133 STs (leaves). This figure appears in color in the electronic version of this article, and any mention of color refers to that version.

### 6.2 Data Results

The proposed approach produces estimated class-specific response profiles 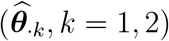 that exhibit differential enrichment of MGEs (Figure 5(b)). For example, MGEs 3, 10 to 17 are estimated with probability of between 0.15 and 0.71 being present in class 1, with log odds ratios 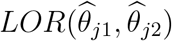 greater than one. The functional annotations of these MGEs reveal that class 1 is likely associated with food-animal hosts. In contrast, MGEs 4 to 9 are estimated to be present in class 2 with probability between 0.35 to 0.82 with LORs greater than one relative to the corresponding estimated response probabilities in class 1. The results suggest the MGEs are highly associated with different types of host-origins.

**Figure 5:**
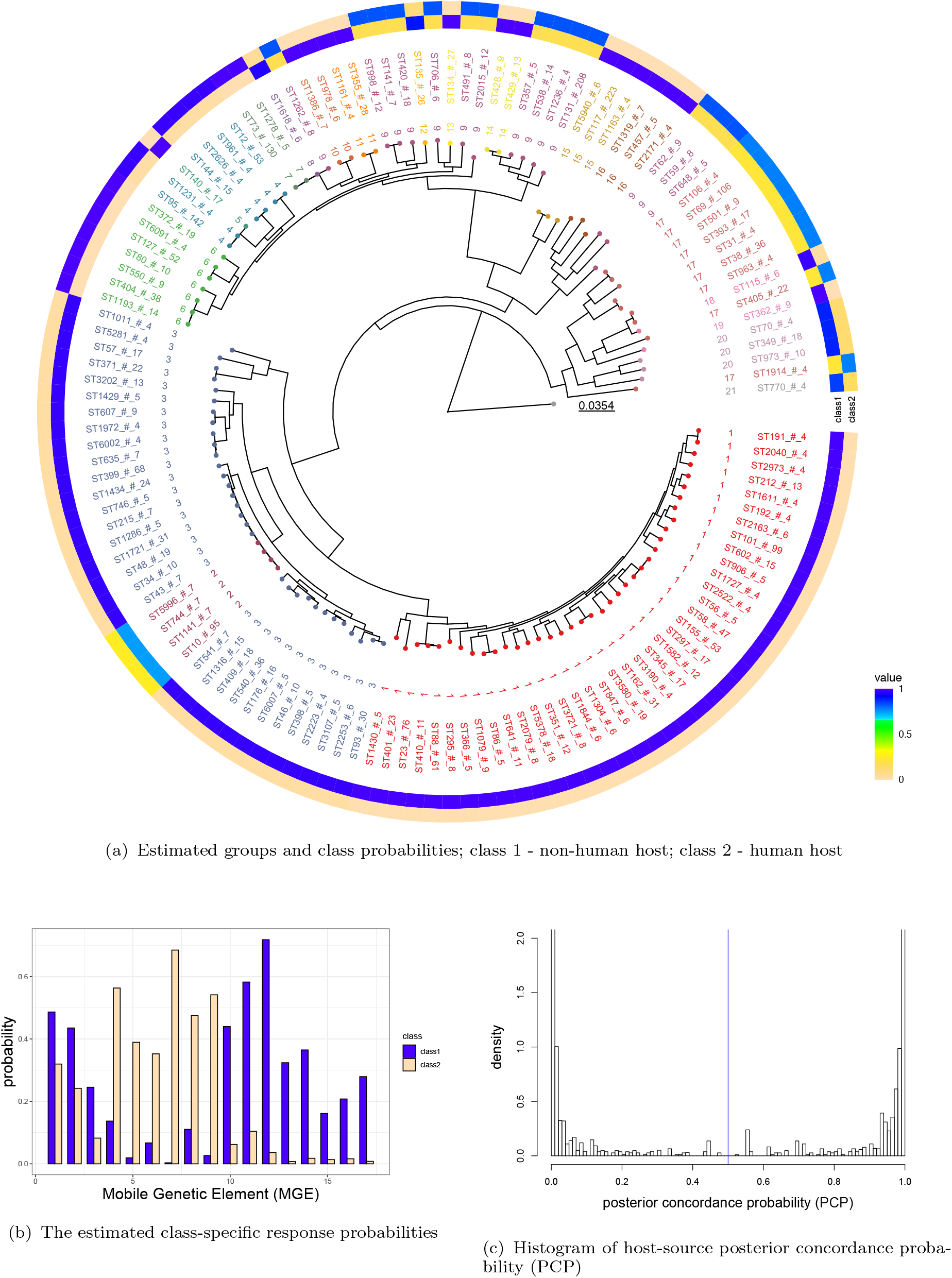
a) Data results with estimated leaf groups and latent class probabilities by group. ST names (ST # isolates) are aligned to the tips of the circular tree, which are colored by discovered leaf groups. The scale bar represents the substitution rate in the conserved core genome. The circular heatmap shows the estimated latent class probabilities 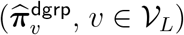; b) and c): see the captions of the subfigures. This figure appears in color in the electronic version of this article, and any mention of color refers to that version.

The proposed approach discovered 21 ST leaf groups, for which distinct estimated vectors of the latent class probabilities 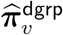 are shown in Figure 5(a). For many estimated ST groups, the class probabilities are almost entirely dominated by one type of host-origin. For example, the estimated ST Group 1 (38 leaves; 649 samples; class 1 probability 0.98, 95% CrI: (0.97, 0.99)) and Group 3 (31 leaves; 422 samples; class 1 probability 0.97, 95% CrI: (0.96, 0.98)) showed high probabilities of non-human (class 1) host-origin of *E. coli*. The results suggest recent cross-species transmissions were rare among multiple nearby lineages. We also compared against results based on two fixed and more restrictive leaf groups, (a) classical LCM (one leaf group); (b) four leaf groups selected by the scientific team (Appendix Figure S4). The single LCM (a) estimated the probability of class 1 to be 0.60, 95% CrI : (0.58, 0.62). The ad hoc leaf grouping (b) produced coarser estimates relative to the proposed 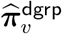 which identified four local leaves (ST1141, ST10, ST744 and ST5996) comprising 116 samples that have estimated probability of class 1: 0.74 (0.66, 0.82)). This highlights the inability of potentially misspecified leaf groups to uncover subtle local variations in the latent class probabilities. We compared these models via 10-fold cross-validation based on the mean predictive log-likelihood (MPL) of the test data, which is computed by plugging in the estimated latent class probabilities and response probability profiles. Of note, because of small sample sizes in some leaves, a naive cross-validation may by chance result in a training set without any observation in some leaves. We therefore randomly keep two observations per leaf and use one random fold of the remaining samples as test data. The proposed approach (with posterior median node selection) achieves the highest MPL (−2015.48) compared to (a) (−2030.15) and (b)(−2162.45). The estimates of response probability profiles are similar. On an individual isolate level, the proposed model can estimate the probability that an isolate was derived from a particular host. For example, by incorporating additional observed sample source information, we can compute “posterior concordance probability (PCP)” for each observation. In particular, PCP, 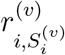, is defined as the approximate posterior probability of the true host origin agreeing with the *observed* sample source category 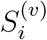 of the same *E. coli* isolate (e.g., 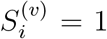 for meat and 2 for human clinical samples). Figure 5(c) shows the histogram of PCPs for all the isolates. Small PCPs, e.g., below a user-specified threshold of 0.5, indicate likely recent host jumps which may subject to further examination to estimate the timing of host transmissions based on *in vitro* stability data of each MGE.

## 7. Discussion

In this paper, we proposed a tree-integrative LCM for analyzing multivariate binary data. We formulated the motivating scientific question in terms of inferring latent class probabilities that may vary in different parts of a tree. We proposed a Gaussian diffusion prior for logistic stick-breaking parameterized latent class probabilities and designed a scalable approximate algorithm for posterior inference. Our *E. coli* data analysis revealed that multiple MGEs are disproportionately associated with specific host origins. Combined with external sample source information, the model can help identify isolates that underwent recent host jump, paving the way for further isolate-level host origin validation.

Our study has some limitations. First, the MGE data we analyzed may represent a fraction of the host-associated accessory elements. By design, additional accessory elements identified in future studies can be readily integrated and evaluated in the proposed framework. Second, host-associated accessory elements are lost and gained over time as *E. coli* strains transition across hosts. For infections that were zoonotic in nature, we did not observe how much time had lapsed between the cross-species host jump and the actual infection. Our model partly accounted for these uncertainties by the imperfect positive response probabilities. However, the timings may drive the presence or absence of multiple MGEs, resulting in potential statistical dependence given the true class of host-origin. Deviations from local independence assumption may impact model-based inference (e.g., Pepe and Janes, 2006; Albert and Dodd, 2004). In practice, a subset of samples with ascertained host-origins may provide critical information to estimate the conditional dependence structure.

Further model extensions may improve model applicability. First, when a subset of observations is not mapped in the tree at random, the algorithm can add additional unobserved leaf indicators to be inferred along with other parameters. Second, it is important to note that the tree integrated into LCM in general is estimated with uncertainty in the topological structure. Methods that use an additional layer of prior over the tree space centered around the estimated tree may account for the upstream uncertainty (e.g., Willis and Bell, 2018). Third, *E. coli* isolates may vary in additional factors such as the hosts’ clinical characteristics. Regression extensions may refine the understanding of variation in latent class probabilities and positive response probabilities that are driven by covariates (e.g., Huang and Bandeen-Roche, 2004). Fourth, LCM is an example of probability tensor decomposition methods (e.g., Johndrow et al., 2017), the tree-integrative LCM motivates extensions to general graph-guided probability tensor decomposition methods. Finally, the truncated stick-breaking formulation in Equation (3) motivates connections to a broader class of covariate-indexed dependent process priors as *K* approaches infinity (e.g., Rodriguez and Dunson, 2011; Ren et al., 2011). Extensions along this line may also relax the present assumption of identical number of realized classes at additional computational cost.

Without relying on prior-likelihood conjugacy, neuronized priors for Bayesian sparse linear regression has been proposed (Shin and Liu, 2021). Comparative studies against spike-and-slab priors are warranted. One known drawback of mean field VI is that it tends to underestimate the marginal posterior variances of parameters. In our simulations, we showed near nominal coverages of the true ***π*** with slight undercoverages happening mostly for leaf groups with very small sample sizes. It is an interesting line of work to incorporate the methods of Giordano et al. (2015) to correct the variance-covariance matrices used in the component variational distributions. We leave these topics for future work.

## Acknowledgment

The research is supported in part by a Precision Health Investigator Award from University of Michigan, Ann Arbor (ML, ZW); an award from Wellcome Trust (LBP, MA and CML; award number 201866); and National Institutes of Health (NIH) grants R01AR073208 (ZW), P30CA04659 (ZW), and 1R01AI130066-01A1 (LBP). We also thank the Co-Editor, Associate Editor and two referees whose comments greatly improved the presentation of our work.

## Data Availability Statement

An R package “lotR” is freely available at https://github.com/zhenkewu/lotR. The data that support the findings in this paper are available from the corresponding author upon reasonable request.

## Supporting Information

Web Appendices and Figures referenced in Sections 4, 5 and 6, and R programs are available with this paper at the *Biometrics* website on Wiley Online Library.

### Algorithm 1

Pseudocode for Variational Algorithm to Integrate Sample Similarities into Latent Class Analysis

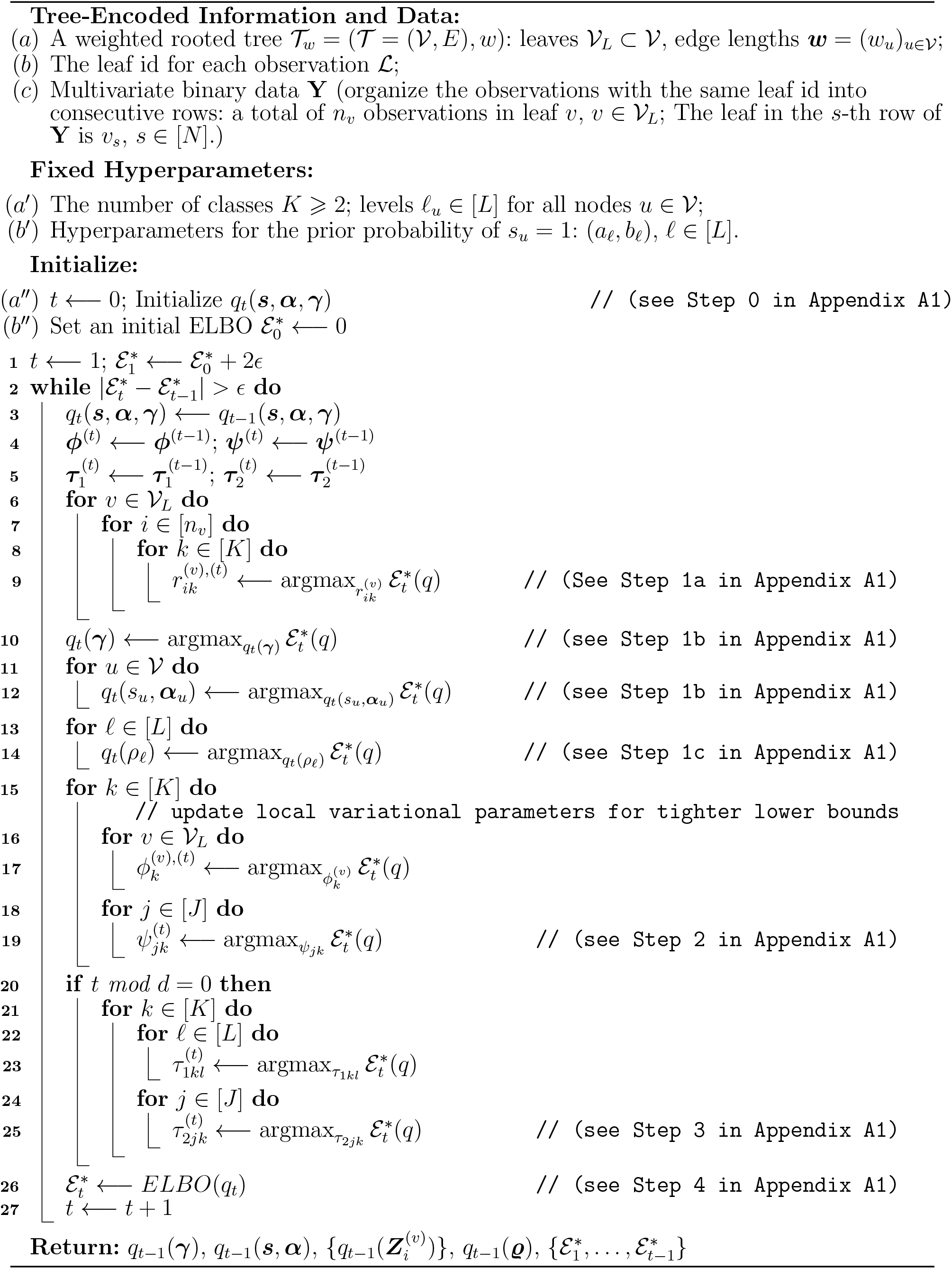

